# Extracellular Electron Transfer Powers *Enterococcus faecalis* Biofilm Metabolism

**DOI:** 10.1101/130146

**Authors:** Damien Keogh, Ling Ning Lam, Lucinda E. Doyle, Artur Matysik, Shruti Pavagadhi, Shivshankar Umashankar, Jennifer L. Dale, Chris B. Boothroyd, Gary M. Dunny, Sanjay Swarup, Rohan B. H. Williams, Enrico Marsili, Kimberly A. Kline

**Author notes:** Equal contribution (Damien Keogh, Ling Ning Lam). **Classification:** Biological Sciences – Microbiology.

## Abstract

Enterococci are important human commensals and significant opportunistic pathogens associated with endocarditis, urinary tract infections, wound and surgical site infections, and medical device associated infections. These infections often become chronic upon the formation of biofilm. The biofilm matrix establishes properties that distinguish this state from free-living bacterial cells and increase tolerance to antimicrobial interventions. The metabolic versatility of the Enterococci is reflected in the diversity and complexity of environments and communities in which they thrive. Understanding metabolic factors governing colonization and persistence in different host niches can reveal factors influencing the transition from commensal to opportunistic pathogen. Here, we report a new form of iron-dependent metabolism for *Enterococcus faecalis* where, in the absence of heme, respiration components can be utilised for extracellular electron transfer (EET). Iron augments *E. faecalis* biofilm growth and generates alterations in biofilm matrix, cell spatial distribution, and biofilm matrix properties. We identify the genes involved in iron-augmented biofilm growth and show that it occurs by promoting EET to iron within biofilm.

**Significance:** Bacterial metabolic versatility is often key in dictating the outcome of host-pathogen interactions, yet determinants of metabolic shifts are difficult to resolve. The bacterial biofilm matrix provides the structural and functional support that distinguishes this state from free-living bacterial cells. Here, we show that the biofilm matrix provides access to resources necessary for metabolism and growth which are otherwise inaccessible in the planktonic state. Our data shows that in the absence of heme, components of *Enterococcus faecalis* respiration (l-lactate dehydrogenase and acetaldehyde dehydrogenase) may function as initiators of EET through the cytoplasmic membrane quinone pool and utilize matrix-associated iron to carry out EET. The presence of iron resources within the biofilm matrix leads to enhanced biofilm growth.

## Introduction

Human colonization by Enterococci initiates immediately at birth via gastrointestinal inoculation from maternal sources, diet, and the environment (1). *Enterococcus faecalis* represents a significant proportion of this early Enterococcal population and remains a stable member of the community throughout life (1). In the gastrointestinal tract, Enterococci are present in the lumen as well as in more specialized niches in the physicochemically complex mucus epithelial layer and epithelial crypts in the small intestine (2).

Bacterial biofilms contribute to host health and disease, and are complex systems that rely on the biofilm matrix to provide the structural and functional properties that distinguish this state from free-living bacterial cells (3). The dynamics of biofilm formation are dependent on many factors such as nutrient availability, environmental stress, social competition, and the generation of extracellular matrix materials. These extracellular polymeric substances provide the architecture surrounding the bacterial cells enabling emergent properties such as resource capture, tolerance to antimicrobial compounds, cooperation or competition, and localized gradients. Infections can become chronic through the development of biofilm. Antimicrobial interventions and colonization by multidrug resistant bacteria can influence endogenous gut microbiome diversity. In both adults and neonates in intensive care settings, gastrointestinal perturbations can result in the replacement of the endogenous Enterococci with aminoglycoside-or vancomycin-resistant Enterococci (4-6). Enterococci are also opportunistic pathogens associated with surgical site and wound infections, bacteraemia, urinary tract infections (UTI), catheter-associated UTI (CAUTI), and endocarditis (7-10). These infections can become chronic through the development of biofilm which is inherently more tolerant to antimicrobial therapies and immune clearance, a problem compounded by multidrug resistant Enterococci in healthcare settings.

The lactic acid bacteria, which include Enterococci, use an electron transport chain for aerobic respiration when external heme is provided or, alternatively, can perform fermentation in the absence of heme. *E. faecalis* does not synthesize heme *de novo* and therefore lacks porphyrin rings required for cytochrome *bd* activity during cellular respiration (11, 12). Similar to other members of the lactic acid bacterial group, *E. faecalis* does not have a major growth requirement for nutritional iron, with manganese instead being used as the essential cofactor for cellular processes (13-16). *E. faecalis* is also remarkably resistant to oxidative stress from O_2_ ^-^, hydroxyl radicals (OH?), and hydrogen peroxide (H O) sources from incomplete reduction of oxygen during respiration; from the oxidative action of host immune cells; or that arise during the Fenton reaction (17, 18). These findings suggest that *E. faecalis* may be able to withstand iron at concentrations which are typically toxic to most bacterial species. In human serum, free iron is far below the 10^-7^ to 10^-5^ M optimal range required to support the growth of most bacteria (19). However, iron availability in the host can vary. Disease states such as haemochromatosis and thalassemia disrupt homeostasis and cause excess iron accumulation (20, 21). Iron oxides have been detected in human spleens and the iron concentration of the gastrointestinal tract is sufficient to support bacterial growth (22, 23). Moreover, *E. faecalis* is tolerant to conditions of significant iron limitation and high oxidative stress (13-18, 24, 25).

In this study, we hypothesize that the ability to withstand higher iron concentrations provides a metabolic advantage to *E. faecalis*. We combine biofilm studies, transmission electron microscopy (TEM), confocal laser scanning microscopy (CLSM), forward genetics, and chronocoulometry to reveal a novel mode of biofilm-specific metabolism in *E. faecalis* where the biofilm matrix harbours iron sinks involved in extracellular electron transport to support biofilm growth. Understanding the metabolic factors that promote colonization and biofilm formation in different environmental conditions may inform mechanisms governing the switch from commensal to opportunistic pathogen and the spread of multidrug resistant Enterococci in a number of ecological reservoirs.

## Results

### Iron supplementation promotes *Enterococcus faecalis* biofilm growth and alters biofilm matrix and matrix properties

To understand the influence of iron on *E. faecalis* biofilm, we evaluated biofilm growth using ferric chloride (FeCl_3_) enriched medium. We analyzed biofilm by confocal laser scanning microscopy (CLSM) in the absence (normal) and presence (supplemented) of additional FeCl_3_ in a flow cell biofilm system. Using a GFP-expressing *E. faecalis* strain, we observed augmented biofilm accumulation as early as 8 hours in medium supplemented with 0.2 mM FeCl_3_ compared to normal medium (Fig. 1a). Iron-enhanced biofilm growth continued for 18 hr, in contrast to the thinner normal biofilm. Iron supplementation did not augment *E. faecalis* growth in planktonic culture (Fig. S1) indicating that the promotion of *E. faecalis* growth is a biofilm specific phenotype. For detailed visualization of *E. faecalis* cell distribution and biofilm architecture properties, we created 3D reconstructions of high magnification CLSM stacks (Fig. 1b). At 18 hr, 37 ^°^C *E. faecalis* biofilm forms a flat monolayer in the normal medium as reported previously (26, 27). By contrast, *E. faecalis* biofilm is thicker and the bacterial cells are distributed throughout the iron supplemented biofilm. Increased biofilm thickness indicates an expansion in biofilm matrix in the supplemented medium suggesting a role for the matrix in iron-enhanced biofilms. We further hypothesized that *E. faecalis* gains a metabolic advantage by the supplementation of iron which in turn affects growth rate. We reasoned that by reducing the temperature we could slow the *E. faecalis* growth rate and possibly modulate the dependence on iron supplementation. At 120 hr, *E. faecalis* grown at 22 ^°^C forms a sparse monolayer biofilm in normal medium; however, the biofilm expands to a depth of greater than 24 µm (the image resolution limit of CSLM) and increased cell density in iron supplemented medium. To evaluate morphological changes in *E. faecalis* cells or the biofilm matrix, we analyzed biofilm cells grown in flow chambers by transmission electron microscopy (TEM). We observed typical diplococcal *E. faecalis* in the normal biofilm cells with an abundance of extracellular fibrous material in the matrix (Fig. 2a,b,c). Constituents of the *E. faecalis* biofilm matrix are not well-understood; however, *E. faecalis* can actively secrete eDNA in biofilms, and DNAse treatment can disperse *E. faecalis* biofilms suggesting eDNA is a major component of the matrix (28-30). The TEM images of the iron supplemented biofilm are similar to normal biofilm (Fig. 2a,b,c), with the exception of electron-dense particles associated with the extracellular fibrous matrix material (Fig. 2d,e,f). Using energy-dispersive X-ray spectroscopy (EDS) in a TEM to analyse the same samples, we demonstrated the extracellular electron-dense particles to be enriched in iron (Fig 2g-m) and hypothesize that these iron deposits together with the increased biomass observed by CSLM are important for augmented *E. faecalis* biofilm formation.

**Figure 1:**
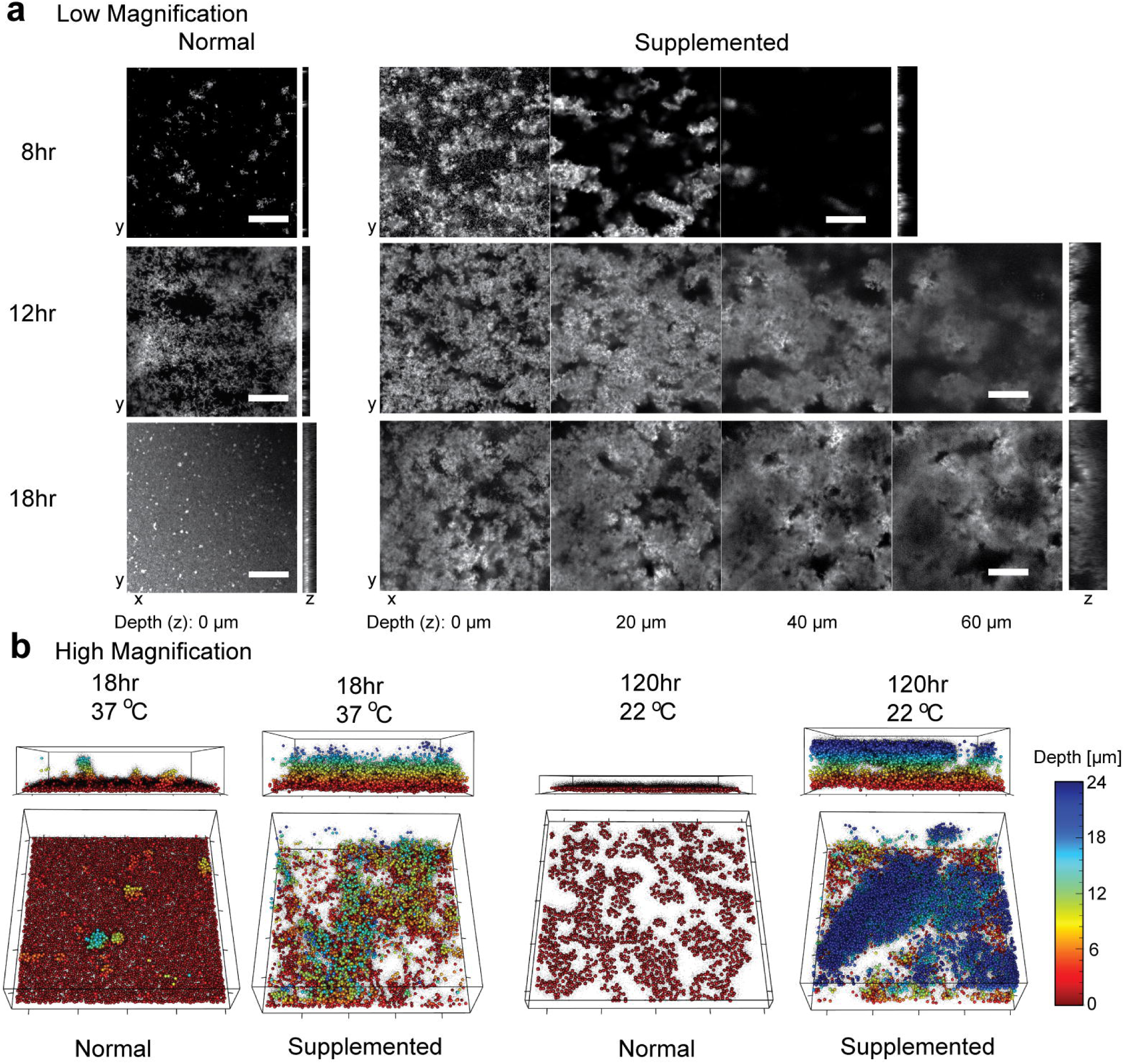
*E. faecalis* flow cell biofilms in iron supplemented media. (**a**) CSLM images at 8, 12, and 18 hr for normal (10 % TSBG) and supplemented (10% TSBG plus 0.2 mM FeCl3). Selected optical sections at the indicated depths are followed by representative lateral view in YZ at the right of each image set. Scale bar 100 μm. (**b**) 3D reconstructions of high magnification CLSM stacks of *E. faecalis* flow cell biofilms grown in normal or supplemented medium for 18 hr at 37 °C or 120 hr at 22 °C. Biofilm depth is color-coded as indicated on z-depth scale (0 to 24 μm) and the lateral box dimensions are 85 × 85 μm (20 μm between ticks).

**Figure 2:**
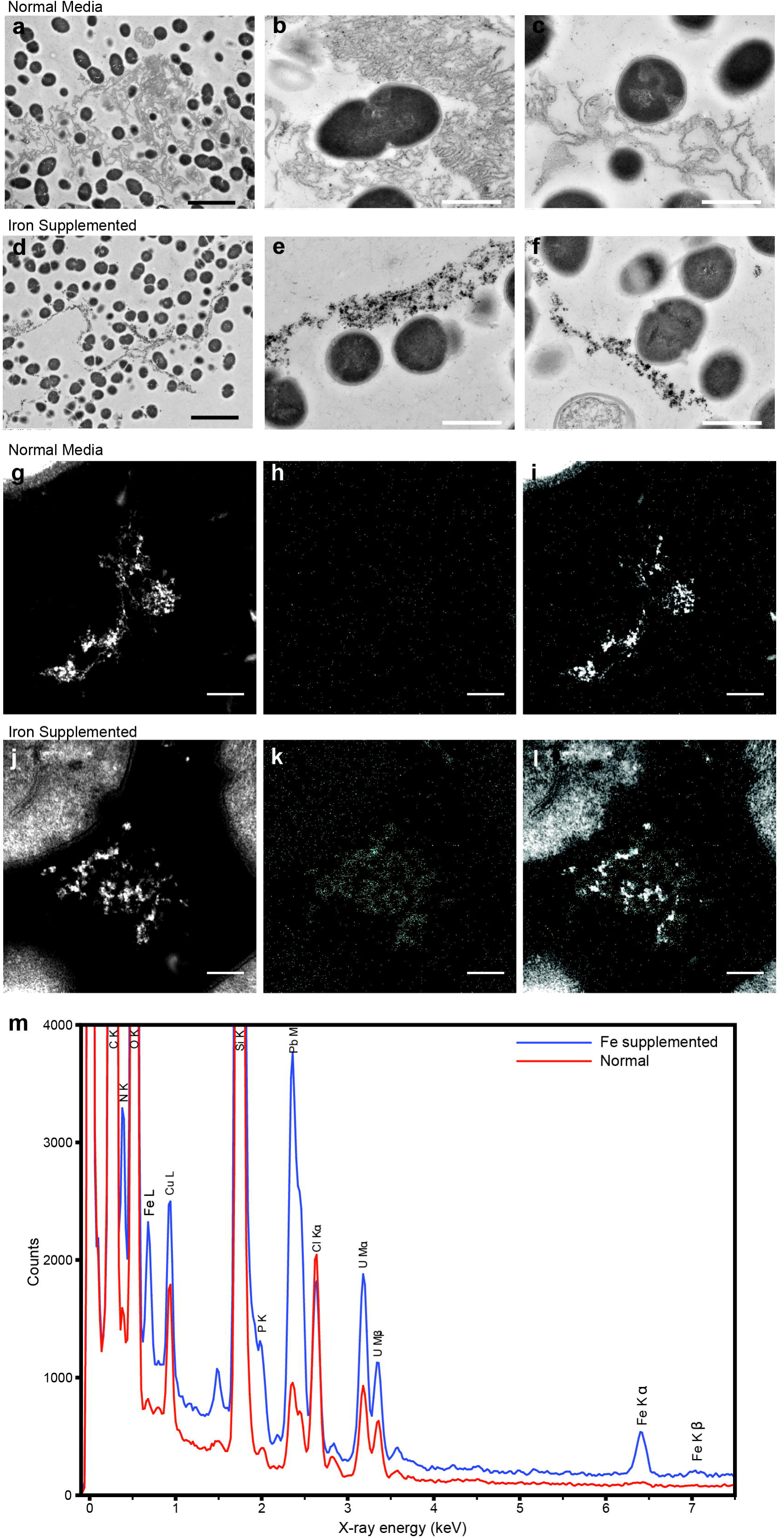
Electron micrographs of the *E. faecalis* biofilm matrix with iron supplementation. Representative images from TEM of *E. faecalis* biofilm from flow cell in normal 10 % TSBG (**a**, **b**, **c**) or 10 % TSBG supplemented with 0.2 mM FeCl3 (**d**, **e**, **f**). In a-f, scale bars represent 2 μm (black) and 0.5 μm (white), and red arrows highlight examples of electron dense particles. The biofilm matrix from biofilms grown in normal media (g, h, i) or with iron supplementation (j, k, l) was examined by HAADF STEM and EDS mapping at 300,000x magnification: HAADF STEM (g, j), iron EDS map (h, k), and merged EDS-STEM images (i, l) are shown at 300,000x magnification. Scale bars in g-l represent 1 μm. The EDS spectra for iron, corresponding to the images are shown in m.

### Iron-induced *Enterococcus faecalis* growth is biofilm specific and iron availability impacts the metallome

To determine if redox-active metals other than iron could also augment *E. faecalis* biofilm formation, we performed biofilm assays with quantification via crystal violet (CV) staining in microtitre plate format to increase throughput (31). In all biofilm assays we included abiotic media controls with metals supplemented to monitor metal precipitation and found that no metals fell out of solution at the concentrations tested in these assays (see Material and Methods – Biofilm Assays, Fig S1). Consistent with *E. faecalis* biofilm augmentation observed when grown in flow cells with 0.2 mM FeCl_3_ supplemented (Fig. 1a,b), FeCl_3_ enriched medium facilitated enhanced *E. faecalis* biofilm growth over time (0 to 120 hr) compared to the normal medium (Fig. 3a). The biofilm increased significantly in response to increasing FeCl_3_ supplementation with 0.5 to 2 mM providing the optimal concentration range in this assay. Above these concentrations, growth was also enhanced but not as significantly compared to the non-supplemented control. We next investigated the capacity of different metal species for enhanced *E. faecalis* biofilm accumulation and found that supplementation with ferrous sulfate (FeSO_4_) or ferric sulfate (Fe_2_(SO_4_)_3_) also increased *E. faecalis* biofilm growth; whereas ferric citrate, magnesium chloride, zinc chloride, or copper chloride had negligible influence on *E. faecalis* biofilm accumulation (Fig 3b). Manganese chloride had a modest augmenting effect on biofilm formation. Heme supplementation at concentrations from 0 to 150 μM also significantly promoted biofilm accumulation and is likely the result of enhanced activity in cytochrome *bd* since heme functions as a cofactor for this enzyme and drives *E. faecalis* to aerobic respiration (32). TEM analysis (Fig. 2d,e,f) revealed iron deposits in the extracellular biofilm matrix of the iron supplemented cultures, whereas no extracellular deposits were observed for biofilms supplemented with heme (Fig S3). We therefore hypothesized that iron concentrations in the *E. faecalis* intracellular metallome would remain stable between the normal and iron supplemented cultures. We used inductively coupled plasma mass spectrometry (ICP-MS) to quantify the intracellular metallome of *E. faecalis* in the absence (normal) and presence (supplemented) of FeCl_3_ supplementation (Fig. 3d, Table S1). The overall intracellular metal content when grown in iron-supplemented medium (3125 ppb) was actually lower than when grown in the normal medium (4641 ppb). Iron was present at 675 ppb in the supplemented samples and 20 ppb in the normal samples. However, the increase in intracellular iron concentration was coincident with a decrease in intracellular cobalt and zinc when *E. faecalis* was grown in iron-supplemented medium. Together, this data suggested that iron was substituting for other metals in the supplemented medium and that the substitution was likely to be functionally balanced. Iron, cobalt, and zinc are transition metals that can function as interchangeable ions and can compete with iron for binding to metalloproteins (33). We hypothesized that identifying genes involved in metal acquisition would help determine if intracellular iron was important for iron-enhanced biofilm growth. We selected 21 mutants of uncharacterized *E. faecalis* membrane transport or regulation systems, with homology to previously characterized metal-associated proteins in other bacteria, from a sequenced mariner transposon library and measured intracellular metal content by ICP-MS (Table S2, S3,S4) (34). These mutants did not have significant changes in intracellular iron concentration in the normal or supplemented medium, with only cobalt and zinc displaying significant changes in some of the mutants and only in the normal medium (Table S2). Because bacterial iron transport systems are generally expressed under iron limitation, we tested these mutants in an iron chelated medium where *E. faecalis* planktonic growth has previously been shown to be growth limited (25). We detected an absence of intracellular iron in 15 mutants under iron limited growth (Table S5) suggesting that these genes govern iron acquisition and may only be active under conditions of low iron availability. Overall, the ICP-MS analysis has demonstrated the modulation of the *E. faecalis* metallome when cultured at different iron concentrations and identified systems involved in metal acquisition. However, this study did not identify any systems involved in iron acquisition in the supplemented medium suggesting that they are not involved or there is a large amount of overlapping functionality in iron transport in *E. faecalis*. These findings support the hypothesis that extracellular electron-dense particles are important for augmented *E. faecalis* biofilm growth in the presence of iron.

**Figure 3:**
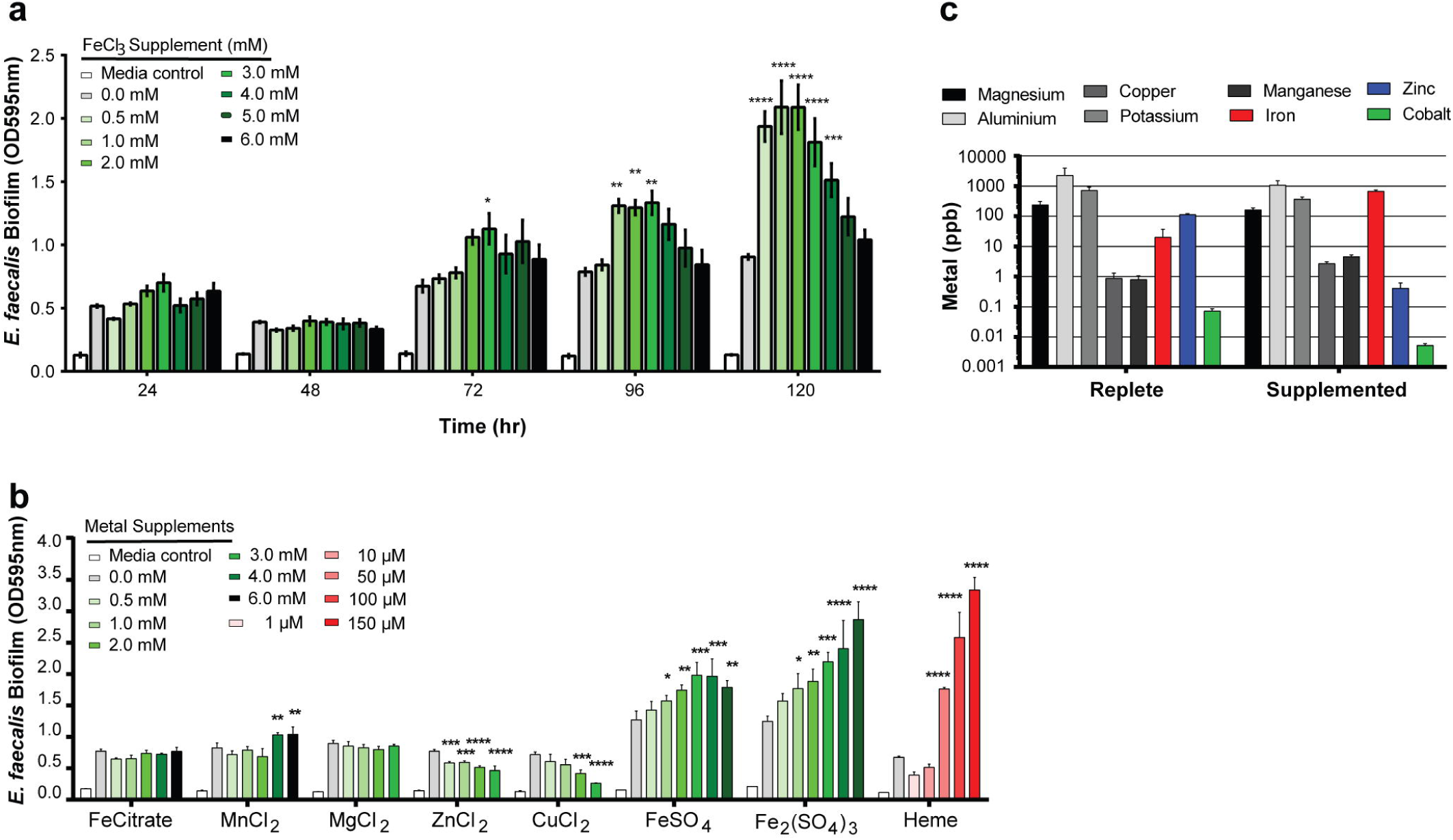
*E. faecalis* biofilm growth under metal supplementation. (**a**) Time course of *E. faecalis* biofilm growth in TSBG supplemented with FeCl3. (**b**) *E. faecalis* biofilm growth at 120hr in TSBG supplemented with metals as indicated. (**c**) ICP-MS analysis of *E. faecalis* grown in TSBG (normal) and TSBG supplemented with 2mM FeCl3 (Supplemented). (**a** and **b**) Data at each time point or metal supplement represents an independent experiment, with the data merged for representation. (**a-c**) n = 3 biological replicates. Statistical significance was determined by two-way ANOVA with Tukey’s test for multiple comparisons, n = 3 with four technical replicates, * *P*=0.05, ** *P*=0.01, *** *P*=0.001, **** *P*=0.0001. (**a**) Statistical analysis was calculated with time set as a ‘repeated measure’. (**a**, **b**, **c**) error bars represent standard deviation (SD) from the mean.

### Components of *E. faecalis* metabolism contribute to iron-induced biofilm growth

To determine the mechanism underlying iron-enhanced *E. faecalis* biofilm, we screened a near-saturated *E. faecalis* mariner transposon library for changes in biofilm formation in media supplemented with 2 mM FeCl_3_ (34). Using the CV biofilm assay, we screened the transposon library for mutants displaying either loss or further enhancement of biofilm growth (Fig. S4a). Mutants displaying altered biofilm growth in iron supplemented media were also examined for alterations in planktonic growth in normal or supplemented media, to eliminate mutants with general fitness defects (Fig. S4a). Because we were seeking factors specifically involved in biofilm formation in excess iron, and not general biofilm factors, we performed a secondary biofilm assay to exclude mutants that displayed altered biofilm in both iron supplemented and normal medium (Fig. S4b-g). The final *E. faecalis* transposon mutants that specifically altered biofilm formation in excess iron included *phoH*, *ldh1*, *trxB2*, OG1RF_11340, OG1RF_10589, and an intergenic region (Table S6). The predicted functions for all of the gene products were associated with metabolism functions and membrane transport.

### Extracellular electron transfer to iron occurs in *E. faecalis* biofilm

Dissimilatory metal-reducing bacteria, such as *Shewanella* spp. and *Geobacter* spp., take advantage of extracellular redox-active metals as terminal electron acceptors for respiration by EET (35, 36). *Shewanella* spp. and *Geobacter* spp. both have functional iron transport and regulation systems for intracellular iron homeostasis (37, 38). EET can function by both direct electron transfer (DET), which involves outer membrane c-type cytochromes (39) and mediated electron transfer (MET), which avails of microbially produced redox mediators (40-43). Our data indicate that genes involved in energy production, redox control, and membrane transport contributed to iron-augmented biofilm formation (Table S2). We also observed iron deposits in the biofilm matrix that surrounds the bacterial cell surfaces of iron-enhanced biofilm (Fig. 2d,e,f,k,m). We therefore hypothesized that the extracellular iron associated with *E. faecalis* biofilm matrix may function as electron sinks via either DET or MET during biofilm metabolism, where Fe(III) is reduced to Fe(II) by *E. faecalis*. To test this, we used chronocoulometry to measure EET of *E. faecalis* biofilms grown in electrochemical cells on a carbon screen-printed electrode (SPE) maintained at high oxidative potential. In the case of DET, extracellular Fe(III) would be reduced by *E. faecalis* and subsequently re-oxidized in part by the electrode in order to regenerate it for further use as an electron acceptor by *E. faecalis*. Alternatively, if the specific mode of EET was MET, we would detect iron acting as a mediator/electron shuttle, carrying electrons extracellularly between *E. faecalis* and the electrode. Electrons are only detected at the electrode surface, and as such any intracellular iron reduction cannot be measured. Thus, in the case of either DET or MET, chronocoulometry allows for the monitoring whether *E. faecalis* is capable of sustained EET when oxidized iron was in its environment in real-time. Additionally, this electrochemical technique measures the integral of current over time, i.e., the charge transferred to the electrode over time (35). Chronocoulometry is commonly used to measure the electron transfer process in redox systems (35, 44). In whole cell bioelectrochemical systems, chronocoulometry is preferred over chronoamperometry to provide a quantitative measure of charge transfer (45). Chronocoulometry was previously used to measure MET in *Pseudomonas aeruginosa* (42). We observed that *E. faecalis* biofilms grew on the carbon electrode (Fig S5a,b) and generated extracellular current in iron supplemented medium, while all abiotic media controls and *E. faecalis* biofilms grown in normal medium did not yield a current (Fig. 4a). Moreover, extracellular current generation was specific to the addition of iron, because we observed no current upon the addition the other biofilm-augmenting metals magnesium, manganese, or heme (Fig. S5c). We then tested the six mutants identified through the transposon screen for the generation of extracellular current in the supplemented medium. Two of the six mutants displayed significantly reduced current as compared to the parental strain. These mutants contained transposon insertions in the genes *ldh1* and OG1RF_11340 suggesting that l-lactate dehydrogenase and acetaldehyde dehydrogenase were involved in extracellular electron transfer in *E. faecalis*. Complementation studies could not be performed because all attempts to clone *ldh* or *adh* on a plasmid *in trans* resulted in an accumulation of mutations preventing the expression of the gene product, presumably due to cytotoxic effects of heterologous metabolic enzyme expression in *E. coli*. To validate that extracellular iron was required as either an electron acceptor (DET) or electron mediator/shuttle (MET), we spiked the electrochemical cell with the chelator 2,2Ldipyridyl to sequester iron. In the presence of iron chelation, we observed a significant reduction in current (Fig. 4b), demonstrating that extracellular iron was instrumental to the process of EET.

**Figure 4:**
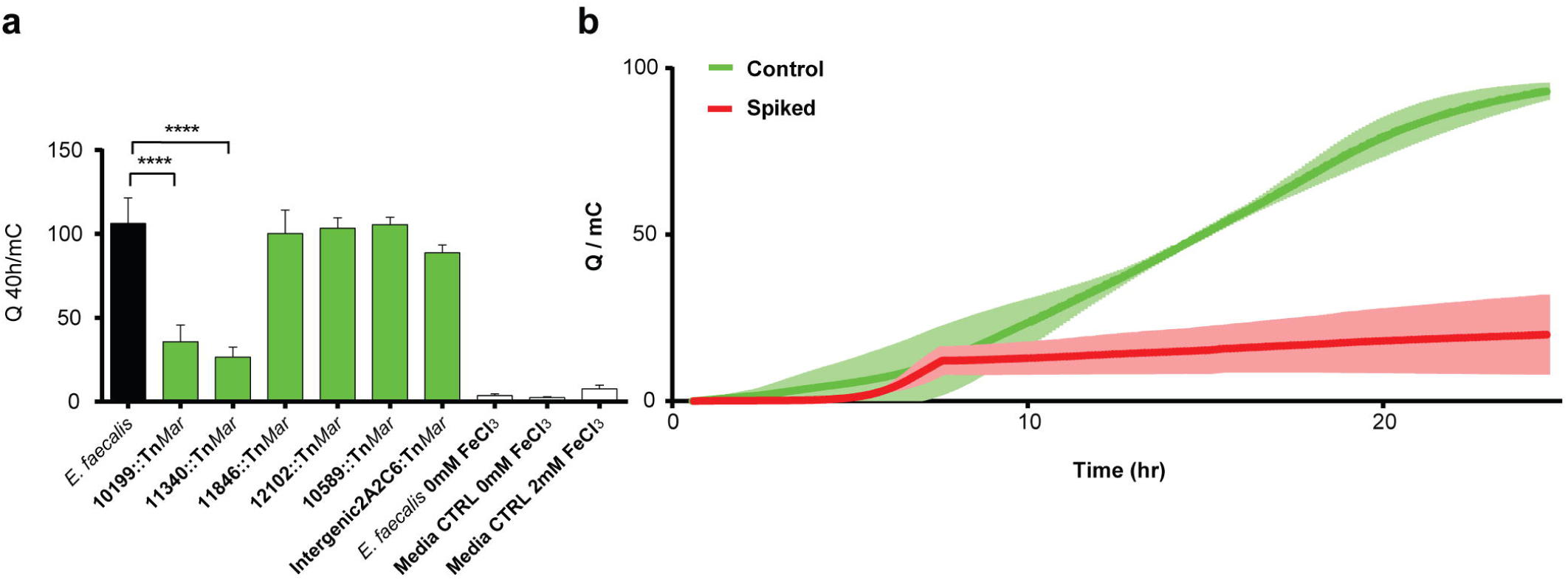
Extracellular electron transfer in *E. faecalis* biofilm. (**a**) Chronocoulometry current (Q) measurement, expressed in milicoulombs (mC), of *E. faecalis* biofilm on a screen-printed carbon mini-electrode over 40 hr in TSBG supplemented with 2mM FeCl3 (Supplemented). Abiotic controls and media controls are indicated. Statistical significance was determined by one-way ANOVA with Tukey’s test for multiple comparisons, n = 3 biological replicates, error bars represent SD from the mean, **** *P*=0.0001. (**b**) Chronocoulometry of *E. faecalis* biofilm in iron supplemented medium with a chelator spike (10 mM 2,2LJdipyridyl) at 7.5 hr. Representative data from four independent experiments are shown, where the trend is consistent among all experiments. Statistical significance was determined by a paired two-tailed t-test, error bars (light-green or-red shading) represent SD from the mean, **** *P*=0.0001.

Taken together, our results establish a model for *E. faecalis* biofilm metabolism where EET using biofilm matrix associated iron supports biofilm growth (Fig. 5). We propose that in the absence of heme, as was the condition in this study, where cytochrome *bd* would be non-functional for aerobic respiration, fermentation end-products such as lactate and acetaldehyde can be used as substrates by l-lactate dehydrogenase (LHD) and acetaldehyde dehydrogenase (ADH) to generate a modification of the electron transport chain. We suggest that these dehydrogenases may act as electron donors transferring electrons to the cytoplasmic membrane quinone (dimethyl menaquinone) pool and ultimately use iron in the biofilm matrix to carry out EET.

**Figure 5:**
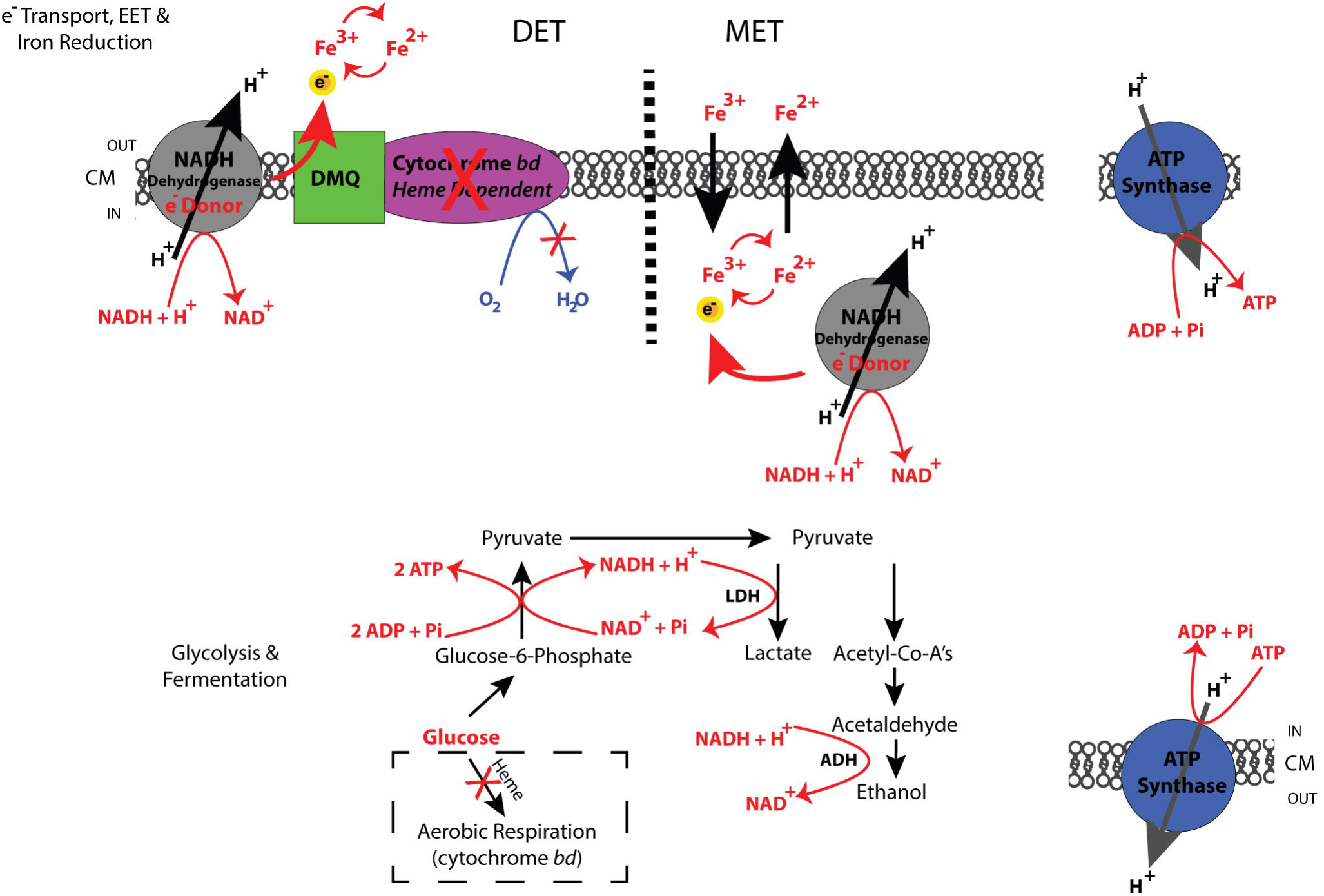
Fermentation and EET-dependent respiration metabolism in *E. faecalis* biofilm. Schematic model for *E. faecalis* biofilm metabolism describing EET through DET and MET mechanisms. ATP is generated by proton flow through membrane integrated ATP synthases. Glycolysis and fermentation are required prior to EET-dependent respiration. This generates the fermentation end-products required as substrates for dehydrogenases acting as electron donors. DET occurs in the absence of heme, where extracellular iron can be reduced and thereby serve as an iron sink for electrons of the respiratory electron transport chain. MET occurs in the absence of heme, when iron is transported intracellularly and reduced, thereafter being exported and serving as an electron mediator/shuttle.

## Discussion

In this study, we characterize a new form of *E. faecalis* metabolism where EET utilising iron results in superior biofilm growth. Understanding the diversity of metabolic possibilities may be important for understanding how *E. faecalis* colonizes different niches such as the human gastrointestinal tract which harbours an abundance of iron (46). Iron rich niches also occur in disease states such as haemochromatosis and thalassemia where iron homeostasis is defective (20, 21). *E. faecalis* is a significant opportunistic pathogen and biofilm development strengthens its tolerance to antimicrobial therapies and immune clearance (7-10). Enterococcal metabolic versatility supports its survival in diverse and complex environments and communities, thus increasing the possibility of dissemination to new niches when iron is present (47). Communities within biofilm are dependent on nutrient and oxygen gradients, cell-to-cell contact and communication, and metabolic cooperation (3). Matrix-associated iron deposits are accessible simultaneously by a number of *E. faecalis* cells and can therefore function as a shared biofilm community resource to enhance biofilm growth.

Our findings demonstrate that *E. faecalis* is electroactive and can utilize EET using iron in the biofilm matrix, via either DET or MET, to enhance biofilm growth. In our model, glycolysis and fermentation proceed prior to EET-dependent respiration to generate the end-product substrates for dehydrogenases acting as electron donors. DET would occur by a mechanism where extracellular iron can be reduced and become an iron sink for electrons of the respiratory electron transport chain. Part of this reduced iron is then reoxidized at the electrode, serving as an electron mediator/shuttle. While further studies are needed to resolve the details of the EET mechanism to iron and to the electrode, these findings extend the current knowledge regarding the extent of EET-capable bacterial species and environments, which are primarily viewed in the context of energy recovery in systems such as microbial fuel cells (36). The characterization of a role for EET in *E. faecalis* metabolism furthers our understanding of possible mechanisms governing colonization and persistence in host niches. Previous studies have demonstrated cell surface-associated oxidation/reduction potential and capacity to utilize exogenously added soluble redox mediators in some Gram-positive bacteria (48, 49). By contrast, we use chronocoulometry to quantify iron-related metabolism in real time, as seen by the sustained generation of an electron flow by the biofilm, leading to augmented biofilm growth. Chronocoulometry quantifies charge transfer at the electrode surface and as such can only detect extracellular iron reduction. In addition, chronocoulometry can show bacterial that derived extracellular electron transfer is sustainable, whereas other commonly used techniques such as cyclic voltammetry can only demonstrate the presence of redox-active moieties on the cell surface and cannot predict whether sustained electron transfer will be possible.

*E. faecalis* and other lactic acid bacteria (LAB) encode the components necessary for aerobic respiration metabolism (50). These are: (1) NADH dehydrogenases functioning as electron donors, (2) a quinone pool to transfer electrons to (3) the terminal electron acceptor complex cytochrome oxidase (51). The *E. faecalis* cytochrome *bd* enzyme can only use oxygen as a terminal electron acceptor when exogenous sources of its co-factor heme are available (11, 52). *E. faecalis* does not synthesise porphyrin and in the absence of heme relies on fermentation. Our experimental assays mimic the absence of heme as this is absent from the growth medium. Our bioelectrochemical data suggests that *E. faecalis* l-lactate and acetaldehyde dehydrogenases function as electron donors for EET to biofilm matrix associated iron sinks. Dehydrogenases transfer hydride from one substrate to another in a reversible reaction that relies on the interconversion of NADH and NAD^+^. Previous studies suggest that, in contrast to most bacterial species that rely on the tricarboxylic acid cycle (TCA) to produce NADH, LAB instead rely on fermentation for its production (53). Fermentation therefore is a necessary metabolic phase prior to the use of EET to support biofilm metabolism. In *Lactococcus lactis*, a fermenting LAB strain, activation of respiration metabolism in the absence of heme and oxygen reduction occurs by copper reduction via the menaquinone pool (54). This supports our finding that extracellular iron functioning as a electron acceptor and/or mediator can activate respiration metabolism in the absence of heme and cytochrome *bd* activity.

Our biofilm data demonstrates an iron-specific *E. faecalis* enhanced biofilm growth response, with only FeCl_3_, heme, FeSO_4_ and Fe_2_(SO_4_)_3_ stimulating the response. Heme availability enables respiration through cytochrome *bd* activity in the presence of oxygen (32). FeSO_4_ and Fe_2_(SO_4_)_3_ are more labile to oxidative changes and precipitation, required for DET-driven iron reduction, than the more soluble ferric citrate which does not enhance biofilm growth. While flow cell biofilm generated enhanced growth at 0.2 mM FeCl_3_, the static biofilm assays only detected enhanced growth from 0.5 to 2 mM FeCl_3_ supplementation.

*E. faecalis* can flourish in iron limited environments and tolerate oxidative stress which can be induced by a high iron environment, contrasting with many other bacterial species where iron is an essential growth nutrient and requires strict iron homeostasis (55). Nonetheless, our ICP-MS analysis shows that iron represents a large proportion of the *E. faecalis* metallome under normal conditions. The modulation of the *E. faecalis* metallome with changes in conditions of either iron limitation or abundance suggests alternative roles for this metal. The absence of detectable iron in the *E. faecalis* mutants with disruptions in genes predicted to encode iron transport and regulatory components, analyzed by ICP-MS under iron limitation, suggests these gene products function in iron acquisition. Previous transcriptomic studies propose roles for *E. faecalis* transport systems in iron acquisition; however our ICP-MS analysis is the first functional characterization of these systems (56, 57). Manganese has been reported to substitute for iron as a cofactor in essential cellular reactions of LAB species, however our data demonstrate that iron represents a greater proportion of the *E. faecalis* metallome than manganese in normal and supplemented conditions (14, 15). The ICP-MS analysis equally highlights that a number of other metals such as zinc, potassium, and magnesium are abundant. Cobalt and zinc are important metals for bacterial growth and we have identified a number of genes governing cobalt and zinc utilization in *E. faecalis* (58). While the intracellular iron levels in the iron supplemented medium was greater than that of the normal condition, no iron transport systems could be identified under these conditions. The absence of detectable transport systems by ICP-MS and the presence of extracellular electron dense deposits in our TEM data suggests that our model of EET would favour DET rather than MET. However, overlapping functionality in nutrient transport is a common strategy for bacteria so single gene mutations may not be sufficient in this condition. Intracellular storage of iron may be strategic in anticipation of acceptor-limited conditions where this metal can be deposited extracellularly.

Colonization and persistence in host niches require adaptability in the exploitation of available resources for energy production and growth. *E. faecalis* is an important human commensal and opportunistic pathogen present in the gastrointestinal tract throughout life. Our work highlights a new form of metabolism where, in the absence of heme, components of respiration can be utilized for EET using extracellular iron. This deeper understanding of mechanisms governing biofilm metabolism in human and clinically relevant bacterial species will enhance approaches to modify or eradicate these reservoirs.

## Materials and Methods

### Bacterial Strains and Growth Conditions

*Enterococcus faecalis* OG1RF (ATCC47077)(59) was grown in Brain Heart Infusion broth (BHI) and cultured at 37 ^°^C under static conditions. *E. faecalis* SD234 is an OG1RF strain derivative harbouring a constitutively expressed *gfp* gene (60). Overnight cultures were normalized to 2-4×10^8^ CFU/ml in phosphate buffered saline (PBS), equivalent to OD_600nm_ 0.7 for *E. faecalis*. For planktonic and biofilm assays, bacteria were cultured at 37 ^°^C (under 200 rpm orbital shaking or static conditions, respectively) with tryptone soya broth supplemented with 10 mM glucose (TSBG) and solidified with 1.5 % agar when appropriate (Oxoid Technical No.3). BHI was supplied by Becton, Dickinson and Company, Franklin Lakes, NJ. TSB and agar was supplied by Oxoid Inc., Ontario, Canada. Metals for supplementation were added during medium preparation. These metals and the chelator 2,2’dipyridyl were supplied by Sigma Aldrich, St Louis, MO, USA.

### Biofilm Assay

Bacterial cultures were normalized as described above and inoculated at 1.6-3.2×10^6^ CFU/200μl microtiter well in TSBG in a 96-well flat bottom transparent microtiter plate (Thermo Scientific, Waltman, MA, USA), and incubated at 37^°^C under static conditions. Uninoculated media controls with metals supplemented to the highest concentration relevant to the assay were included to check for supplemented metal precipitation. Supernatants were discarded and the microtitre plate washed twice with PBS. To stain surface adherent bacteria, 200 µl of crystal violet solution at 0.1% w/v (Sigma-Aldrich, St Louis, MO, USA) was added to each well and incubated at 4 ^°^C for 30 minutes. This solution was discarded and the microtitre plate washed twice with PBS followed by crystal violet solubilization with 200 µl per well ethanol-acetone (4:1) for 45 minutes at room temperature. The intensity of crystal violet staining was measured by absorbance at OD_595nm_ using a Tecan Infinite 200 PRO spectrophotometer (Tecan Group Ltd., Männedorf, Switzerland).

### Flow Cell Biofilm Assay

Flow cell biofilm studies were performed as previously described with minor modifications (61). Bacterial cultures were normalized to 2-4×10^6^ CFU/ml in PBS and 250 µl of this stock injected through the Stovall flow cell system inlet silicon tube connected to the flow cell chamber. This inoculation was performed when the system flow was halted by clamping both the inlet and outlet silicon tubes. The chamber was inverted for one hour to facilitate bacterial adherence to the glass slide surface. The flow cell system was then reset, unclamped, and the medium feed was set to 4.5 ml/hr.

### Transposon Library Screen

The cryogenically stocked, 96-well format *E. faecalis* OG1RF mariner transposon library consisted of 14978 individual mutants (34). This library was cultured using a cryo-replicator (Adolf Kühner AG) to inoculate DeepWell blocks (Greiner Bio-One) containing 1ml BHI medium for overnight incubation at 37 ^°^C with shaking at 220 rpm. Cultures were normalized to OD_600nm_ 0.1 (2-4×10^8^ CFU/ml) in PBS with the Tecan Infinite**®** 200 PRO spectrophotometer (Tecan Group Ltd., Switzerland) using a 96 well microtiter plate. The primary screen of the library was performed by inoculating a microtitre well with 1.6-3.2×10^6^ CFU/200μl in TSBG medium supplemented with 2 mM FeCl_3_. The microtitre plates were then incubated at 37 °C, statically, inside a moistened chamber to prevent evaporation of media. Biofilm was quantified by crystal violet staining as described above. Mutants with either reduction or further enhancement of biofilm signal compared to wild type controls were then validated using two independent biological replicates for each mutant in TSBG biofilm assays. This primary validation was followed by a planktonic growth validation in TSBG and TSBG medium supplemented with 2 mM FeCl_3_. A secondary validation using three independent biological replicates for each mutant was performed in biofilm assays with TSBG and TSBG medium supplemented with 2 mM FeCl_3_ to eliminate any mutants exhibiting defects in biofilm formation under normal conditions.

### Mapping Tn Insertions

gDNA was extracted using the Wizard**®** Genomic DNA Purification Kit (Promega) from transposon mutants that were not originally mapped. The gDNA quantified and assessed for nucleic acid quality by Qubit High Sensitive dsDNA assay (Invitrogen) and NanoDrop. Sequencing was performed using an Illumina MiSeq. *de novo* reads were assembled using CLC Genomics Workbench version 8.0 and *E. faecalis* OG1RF as a template. The Tn insertion site was identified by BLAST using the mariner transposon sequence and identification of the flanking genomic sequence. Transposon mutant strains were named to include library location information, gene name or intergenic information, followed by “TnMar”. For example, (4.2A1 F1)10589:TnMar indicates the transposon mutant for OG1RF_10589 located in block 4.2A1 of the library in position F1 of the microtitre plate.

### Electrochemical Setup and Analysis

Screen printed electrodes (SPE) (model DRP-C110; DropSens, Spain) consisting of a carbon working electrode, carbon counter electrode, and silver reference electrode were controlled by a multichannel potentiostat (Bio-Logic, France) in an electrochemical cell of 9 ml working volume sealed with a Teflon cap. Chronocoulometry was used to characterize the electrochemical activity of live microbial cultures by measuring the charge passed over the course of growth, with the total charge in mC at 40 hr used for comparison. During chronocoulometry, the working electrode was poised at 200 mV vs the silver pseudoreference electrode of the SPE. This potential was chosen as it is high enough to re-oxidise the reduced iron, while being low enough to avoid damaging the bacteria. By applying a set potential to an electrode acting as an electron acceptor, the extracellular current generated can be tracked with time, the integral of which represents the total charge passed. By looking at the charge passed at a fixed time, the electroactivity of a culture/mutant etc. can be established. Bacterial stocks of 2-4×10^8^ CFU/ml for electrochemical experiments were prepared as described above, and electrochemical cells inoculated to 2-4×10^5^ CFU/ml. All electrochemical experiments were conducted at 37 ^°^C using TSBG medium supplemented with 2 mM FeCl_3_ unless otherwise stated. The iron chelator, 2,2Ldipyridyl (Sigma Aldrich, USA), was spiked in selected experiments to quench EET.

### Thin-section Transmission Electron Microscopy (TEM)

Biofilms were grown using the flow cell protocol or the standard biofilm protocol described, but with the latter using a 6-well plate. The flow cell biofilm was resuspended in a 2 % paraformaldehyde-2.5 % glutaraldehyde solution (Polysciences Inc., Warrington, PA) in 100 mM PBS (pH 7.4) for 1 hr at room temperature. The samples were then embedded in 2 % low-melt agarose, washed in PBS and post-fixed in 1 % osmium tetroxide for 1 hr. Samples were rinsed extensively in distilled water (dH_2_O) prior to *en bloc* staining with 1 % aqueous uranyl acetate (Ted Pella, Inc., Redding, CA, USA) for 1 hr. Following several rinses in dH_2_O, samples were dehydrated in a graded series of ethanol and embedded in Eponate 12 resin (Ted Pella, Inc., Redding, CA, USA). Sections of 95 nm were cut with a Leica Ultracut UCT ultramicrotome (Leica Microsystems, Inc., Bannockburn, IL, USA), stained with uranyl acetate and lead citrate, and viewed on a JEOL 1200 EX transmission electron microscope (JEOL USA, Inc., Peabody, MA, USA).

## Energy-dispersive X-ray spectroscopy (EDS) TEM

Samples were prepared as described above for TEM and viewed on an aberration-corrected JEOL ARM 200 cold field-emission gun transmission electron microscope in high-angle annular dark-field scanning transmission (HAADF STEM) mode using spot size 6. The sample was held in a JEOL Be double tilt holder tilted by about 12 ° towards the X-ray detector. STEM images were collected using a JEOL annular dark-field detector on a Gatan Digiscan with 2048×2048 pixels and a dwell time of 10 μs. Energy dispersive X-ray maps were collected using an Oxford Aztec system with a 0.7 sr collection solid angle by scanning the beam with multiple frames over a period of about 5 minutes at a resolution of 512×512 pixels.

### Confocal Laser Scanning Microscopy and 3D Reconstruction

Biofilm morphology, biovolume, and cell distribution were analyzed by CLSM directly from flow cell chamber glass microscope slides at three separate locations (inlet, middle and outlet areas) with 3 individual Z-stack images per technical replicate. Images were acquired using LSM780 confocal microscope (Zeiss, Germany) equipped with 20x/0.8 Plan-Apochromat objective and controlled by ZEN software. Samples were illuminated with a 488 nm Argon laser line and the GFP emitted fluorescence was collected in the 507-535 nm range. Optical sections (425×425 μm) were collected every 5 μm through the entire biofilm thickness and signal from each section was averaged 2-4 times. Fiji software (62) was used for further processing (levels adjustment, stack reslice). For the 3D biofilm reconstructions, optical slices (85×85 μm) were acquired with 63x/1.4 Plan-Apochromat oil immersion lens every 0.3μm through entire biofilm thickness or until loss of the fluorescence signal (due to light scattering, absorption, and possible fluorescence quenching of thick iron-supplemented biofilms). The center of mass for each cell in 3D space was found using MosaicSuite for Fiji (63), then coordinates were filtered in R (64). To visualize the biofilm matrix and spatial organization coordinates were plotted as spheres with cell-size diameter and color coded Z-depth.

### Inductively Coupled Plasma Mass Spectroscopy (ICP-MS)

*E. faecalis* cultures were prepared as previously described with minor modifications (65). Overnight cultures were normalized as described above and 1-2×10^5^ CFU/well was inoculated into DeepWell blocks (Greiner Bio-One) containing 2 ml TSBG medium and incubated overnight at 37 ^°^C under static conditions. Three biological replicates with five technical replicates were prepared for each *E. faecalis* strain and, following incubation, the technical replicates were pooled prior to preparation for ICP-MS. Harvested cell pellets were washed with 10 mM EDTA (Ambion^TM^ Thermofisher Scientific, USA) prepared in LC-MS grade dH_2_O (Sigma Aldrich, St Louis, MO, USA) and washed three times with LC-MS grade dH_2_0. Cells were then concentrated to 2-4×10^8^ CFU/ml in LC-MS grade dH_2_O. Each cell pellet was digested in 500 μl of 69 % nitric acid and 250 μl of 31 % hydrogen peroxide. Following sample digestion, all the samples were diluted with LC-MS grade dH_2_0 to a 2 % (w/v) nitric acid concentration in the final solution.

An Agilent 7700 series model ICP-MS system (Agilent, Santa Clara, CA, USA) was used for simultaneous determination of selected elements [Mg, Al, P, K, Ca, V, Cr, Mn, Fe, Co, Ni, Cu, Zn] in prepared *E. faecalis* samples. Prior to sample measurement and quantification, stock solutions of a multi-element calibration standard (Inorganic Ventures, VA, USA) were serially diluted (0 µg/L to 1000 µg/L) and run on the system. For each measurement (standards, samples, blanks, and quality controls), addition of internal standard (Sc; 100 mg/L, Agilent, USA) was performed to correct for physical matrix effects. Blanks were determined together with samples for every run and the mean of three runs was determined for each sample. Full quantitative analysis was performed against calibration standards for each element. Quality control samples (multi-element calibration solution; 100 µL) were inserted and run at regular intervals during the experiment to ensure reliability of the data and to eliminate signal drift or interference.

For statistical analysis, 10 metals were measured for 22 phenotypes (*E. faecalis* OG1RF wild type and 21 mutants) in three nutrient conditions, with 3 biological replicates for each mutant and in each condition. During the ICP-MS run, 3 technical replicates were measured from each sample, which were then averaged and used for statistical analysis. The concentration of the metals in a blank control (500 μl of 69% nitric acid and 250 μl of 31% hydrogen peroxide) was subtracted from the concentration of metals in the samples. Metals whose levels were below the detection limit were coded as not detected. For statistical analysis, metal concentrations were normalized using log base 10 transformation. To test differences between the metal levels in mutants and the control in each nutrient condition, Welch’s two sample t-tests were performed in R using the t.test function, with correction for multiple comparisons made using the Benjamini-Hochberg procedure.

## Author Contributions

D.K. conceptualized the study. D.K., L.N.L., E.M., and K.A.K. designed the experiments, analyzed data, and prepared the manuscript. D.K. and L.N.L. performed biofilm experiments and analyzed data. A.M. analyzed confocal data and generated 3D reconstruction models. D.K. and S.P. performed the ICP-MS experiments. S.U. and R.B.H.W. analyzed metabolomics data. D.K., L.E.D., L.N.L., and E.M. performed the electrochemistry experiments and analyzed data. L.N.L and C.B. performed EDS TEM. J.D. and G.D. provided the transposon library. All authors reviewed the manuscript.

## Acknowledgments

This work was supported by the National Research Foundation and Ministry of Education Singapore under its Research Centre of Excellence Programme, by the National Research Foundation under its Singapore NRF Fellowship programme (NRF-NRFF2011-11), and by the Ministry of Education Singapore under its Tier 2 programme (MOE2014-T2-2-124). A.M. and his contributions were supported by the National Medical Research Council under its Clinical Basic Research Grant (NMRC/CBRG/0086/2015) awarded to K.A.K. We thank Kenneth Beckman (University of Minnesota) and colleagues for sequencing of the *E. faecalis* transposon library, and Wandy Beatty (Washington University in St. Louis) for performing TEM. We thank SCELSE members Sumitra Debina Mitra, Irina Afonina, Shu Sin Chng, Hans-Kurt Fleming, Scott Rice, and Staffan Kjelleberg, as well as Jeff Gralnick (University of Minnesota) for their critical assessment of the manuscript. The transmission electron microscopy was performed at the Facility for Analysis, Characterization, Testing and Simulation (FACTS), Nanyang Technological University, Singapore.

---

**Figure S1: *E. faecalis* planktonic growth under metal supplementation.** (**a**) Time course enumeration of *E. faecalis* planktonic growth over 24hr in TSBG and TSBG supplemented with 2mM FeCl3. n = 3 biological replicates, error bars represent standard deviation (SD) from the mean.

**Figure S2: Metal precipitation control.** Static incubation of TSBG medium supplemented with FeCl3 over time. n = 2 replicates, error bars represent standard deviation (SD) from the mean.

**Figure S3: Electron micrographs of the *E. faecalis* biofilm matrix in the absence or presence of metal supplementation.** Representative images from TEM of *E. faecalis* biofilm from 24 hr static biofilms grown in normal TSBG (a, e), TSBG supplemented with 2 mM FeCl3 (b, e), or 50 uM heme (c, f). Scale bars represent 2 μm (black) and 0.5 μm (white).

**Figure S4: Transposon library screen for mutants exhibiting changes in biofilm accumulation under iron supplementation.** (**a**) Flow chart of screen process including evaluation criteria for biofilm assays. (**b-g**) Secondary validation data for mutants passing primary evaluation. Light grey indicates normal medium and dark grey indicates iron supplemented medium. Secondary validation bar charts represent individual experiments and are separately presented, rather than being pooled, for greater clarity.

**Figure S5: Biofilm biovolume and metal-dependence of *E. faecalis* EET.** CLSM images of *E. faecalis* biofilm on screen-printed electrode (SPE) at the end of chronocoulometry measurements after 20 hrs growth in (**a**) normal TSBG or (**b**) TSBG supplemented with 2 mM FeCl3. (**c**) Charge (mC) transferred from *E. faecalis* biofilm to the SPE after 20 hr in normal TSBG, TSBG supplemented with 2mM FeCl3, 2mM manganese chloride (MnCl2), 2 mM magnesium sulfate (MgSO4), or 50uM heme. These results were measured in glass electrochemical cells willed with 11 mL growth medium. All the other parameters are the same described in the Materials and Methods section. The working electrode was poised at 200 mV vs the silver pseudoreference electrode of the SPE.

**Table S1: Metallome of *E. faecalis* in normal and iron supplemented conditions**

**Table S2: Metallome comparison of *E. faecalis* mutants versus wild type in normal condition**

**Table S3: Metallome comparison of *E. faecalis* mutants versus wild type in deplete condition**

**Table S4: Metallome comparison of *E. faecalis* mutants versus wild type in supplemented condition**

**Table S5: *E. faecalis* mutants in deplete condition where iron was not detected**

**Table S6: *E. faecalis* components involved in iron-induced biofilm growth**

